# Characterization of *Mollivirus kamchatka*, the first modern representative of the proposed *Molliviridae* family of giant viruses

**DOI:** 10.1101/844274

**Authors:** Eugene Christo-Foroux, Jean-Marie Alempic, Audrey Lartigue, Sebastien Santini, Karine Labadie, Matthieu Legendre, Chantal Abergel, Jean-Michel Claverie

## Abstract

Microbes trapped in permanently frozen paleosoils (permafrost) are the focus of increasing researches in the context of global warming. Our previous investigations led to the discovery and reactivation of two Acanthamoeba-infecting giant viruses, *Mollivirus sibericum* and *Pithovirus sibericum* from a 30,000-year old permafrost layer. While several modern pithovirus strains have since been isolated, no contemporary mollivirus relative was found. We now describe *Mollivirus kamchatka*, a close relative to *M. sibericum*, isolated from surface soil sampled on the bank of the Kronotsky river in Kamchatka. This discovery confirms that molliviruses have not gone extinct and are at least present in a distant subarctic continental location. This modern isolate exhibits a nucleo-cytoplasmic replication cycle identical to that of *M. sibericum*. Its spherical particle (0.6-μm in diameter) encloses a 648-kb GC-rich double stranded DNA genome coding for 480 proteins of which 61 % are unique to these two molliviruses. The 461 homologous proteins are highly conserved (92 % identical residues in average) despite the presumed stasis of *M. sibericum* for the last 30,000 years. Selection pressure analyses show that most of these proteins contribute to the virus fitness. The comparison of these first two molliviruses clarify their evolutionary relationship with the pandoraviruses, supporting their provisional classification in a distinct family, the *Molliviridae*, pending the eventual discovery of intermediary missing links better demonstrating their common ancestry.

**Importance:** Virology has long been viewed through the prism of human, cattle or plant diseases leading to a largely incomplete picture of the viral world. The serendipitous discovery of the first giant virus visible under light microscopy (i.e., >0.3μm in diameter), mimivirus, opened a new era of environmental virology, now incorporating protozoan-infecting viruses. Planet-wide isolation studies and metagenomes analyses have shown the presence of giant viruses in most terrestrial and aquatic environments including upper Pleistocene frozen soils. Those systematic surveys have led authors to propose several new distinct families, including the *Mimivirida*e, *Marseilleviridae, Faustoviridae, Pandoraviridae*, and *Pithoviridae*. We now propose to introduce one additional family, the *Molliviridae*, following the description of *M. kamchatka*, the first modern relative of *M. sibericum*, previously isolated from 30,000-year old arctic permafrost.

## Introduction

Studies started about 30 years ago, have provided multiple evidence that soils frozen since the late Pleistocene and predominantly located in arctic and subarctic Siberia, do contain a wide diversity of microbes that can be revived upon thawing (1-3) after tens of thousands of years. These studies culminated by the regeneration of a plant from 30,000 year-old fruit tissue (4). Inspired by those studies, we then isolated from a similar sample two different Acanthamoeba-infecting large DNA viruses, named *Pithovirus sibericum* and *Mollivirus sibericum*, demonstrating the ability for these viruses, and maybe many others, to remain infectious after similarly long periods of stasis in permafrost (5-6). Several modern relatives to the prototype Pithovirus have since been isolated and characterized, leading to the emergence of the new proposed *Pithoviridae* family (7-10). In contrast, and despite the increasing sampling efforts deployed by several laboratories, no other relative of *M. sibericum* was found. Without additional isolates, its classification as a prototype of a new family, or as a distant relative of the pandoraviruses (with which it shared several morphological features and 16% of its gene content) (6) remained an open question. Here we report the discovery and detailed characterization of the first modern *M. sibericum* relative, named *Mollivirus kamchatka*, after the location of the Kronotsky river bank where it was retrieved. The comparative analysis of these first two molliviruses highlights their evolutionary processes and suggests their provisional classification into their own family, the *Molliviridae*, distinct from the *Pandoraviridae*, pending the eventual discovery of intermediary missing links clearly establishing their common ancestry.

## Results

### Virus isolation

The original sample consisted of about 50 ml of vegetation-free superficial soil scooped (in sterile tubes) from the bank of the Kronotsky river (coordinates: N 54°32’59’’ E 160°34’55’’) on july 6^th^, 2017. Before being stored at 4°C, the sample was transported in a backpack for a week at ambient temperature (5° C up to 24° C). This area corresponds to a continental subarctic climate: very cold winters, and short, cool to mild summers, low humidity and little precipitation. Back in the laboratory, few grams of the sample were used in the Acanthamoeba co-cultivation procedure previously described (6). After a succession of passages and enrichment on *Acanthamoeba castellanii* cultures, viral particles were produced in sufficient quantity to be recovered and purified.

### Virion morphology and ultrastructure

As for *M. sibericum*, light microscopy of infected cultures showed the multiplication of particles. Using transmission electron microscopy (TEM), these particles - undistinguishable from that of *M. sibericum* - appear approximately spherical, 600 nm in diameter, lined by an internal lipid membrane and enclosed in a 20 nm-thick electron-dense thick tegument covered with a mesh of fibers (Fig. 1).

**Fig 1.**
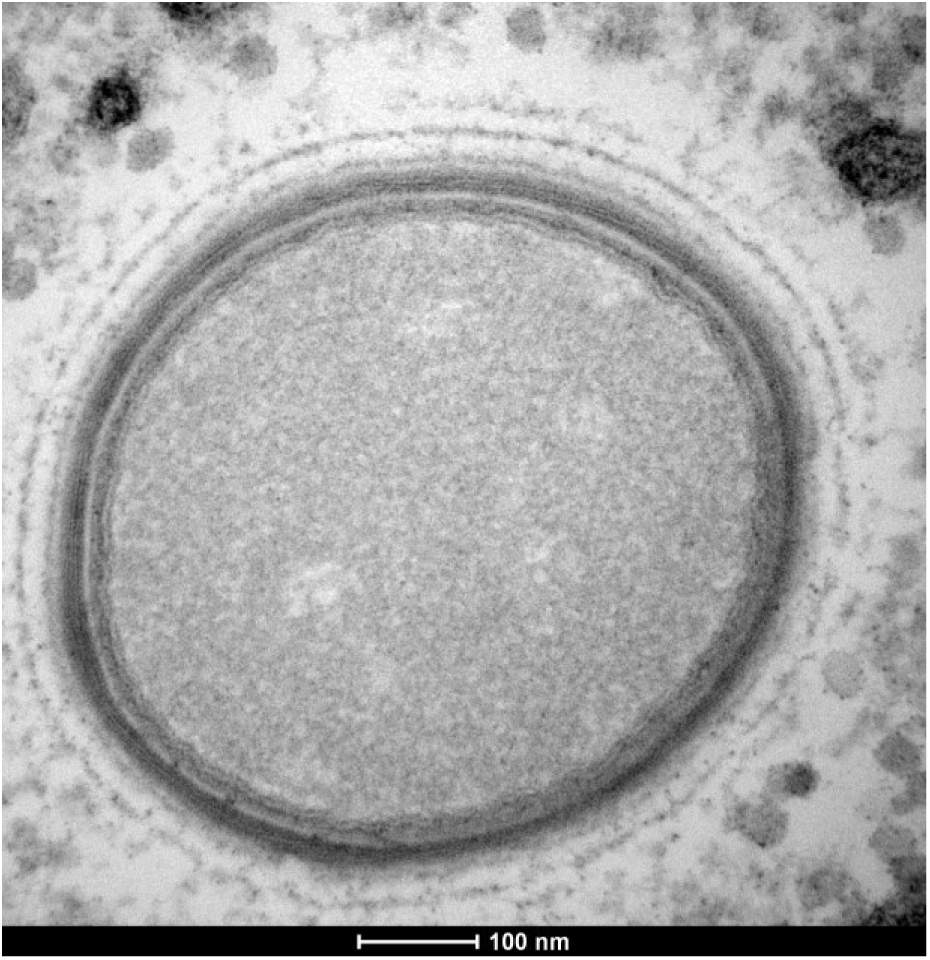
Ultrathin section TEM image of a neo-synthetized *M. kamchatka* particle in the cell cytoplasm 7h post infection. The structure of the mature particles appear identical to that of *M. sibericum.*

### Analysis of the replication cycle

The replication cycle in *A. castellanii* cells was monitored using TEM and light microscopy of DAPI-stained infected cultures as previously described (6). The suite of events previously described for host cells infected by *M. sibericum*, was similarly observed upon *M. kamchatka* replication (6, 11). After entering the amoeba cell through phagocytosis, *M. Kamchatka* virions are found gathered in large vacuoles individually or in groups of 2-6 particles. Multiple nuclear events occur during the infection starting with the drift of the host cell nucleolus to the periphery of the nucleus 4-5h post infection (p.i.). 7h p.i., the nucleus appears filled with numerous fibrils that may correspond to viral genomes tightly packed in DNA-protein complexes (Fig. 2A). About 30% of the nuclei observed at that time exhibit a ruptured nuclear membrane (Fig. 2B). Besides those internal nuclear events, we observed a loss of vacuolization within the host cell from 4h p.i. to the end of the cycle, in average 9h p.i.. Large viral factories are formed in the cytoplasm at the periphery of the disorganized nucleus (Fig. 2). These viral factories displays the same characteristics as those formed during *M. sibericum* infections, involving an active recycling of membrane fragments (11).

**Fig 2.**
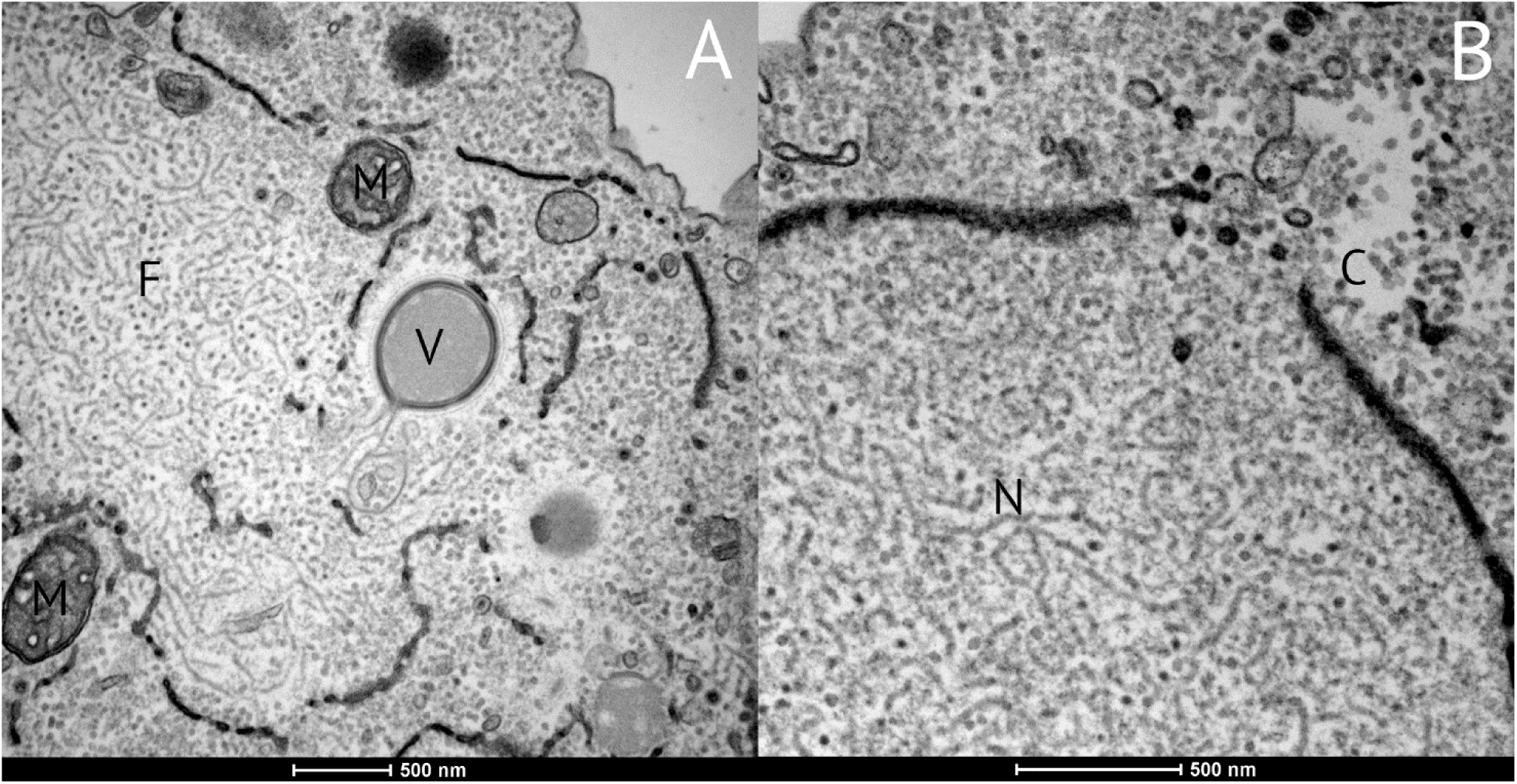
Ultrathin section TEM image of *A. castellanii* cell 7 to 10 hours post infection by *M. kamchatka*. (A) Viral factory exhibiting fibrils (F), a nascent viral particle (V), and surrounding mitochondria (M). Fragments of the ruptured nuclear membrane are visible as dark bead strings. (B) Details of a nuclear membrane rupture through which fibrils synthetized in the nucleus (N) are shed into the cytoplasm (C).

### Comparative genomics

DNA prepared from purified *M. kamchatka* particles was sequenced using both Illumina and Oxford Nanopore platforms. *M. kamchatka* genome, a linear double stranded DNA molecule (dsDNA), was readily assembled as a unique sequence of 648,864 bp. The read coverage was uniform throughout the entire genome except for a 10 kb terminal segment presumably repeated at both ends and exhibiting twice the average value. The *M. kamchatka* genome is thus topologically identical to that of a *M. sibericum*, slightly larger in size (when including both terminal repeats), and with the same global nucleotide composition (G+C= 60%). This similarity was confirmed by the detailed comparison of their genome sequences exhibiting a global collinearity solely interrupted by a few insertions and deletions.

Prior to the comparison of their gene contents, *M. sibericum* and *M. kamchatka* were both annotated using the same stringent procedure that we previously developed to correct for gene overpredictions suspected to occur in G+C rich sequences such as those of pandoraviruses (12-14). A total of 495 and 480 genes were predicted for *M. sibericum* and *M. kamchatka*, with the encoded proteins ranging from 51 to 2171 residues and from 57 to 2176 residues, respectively. *M. kamchatka* predicted protein sequences were used in similarity search against the non-redundant protein sequence database (15) and the re-annotated *M. sibericum* predicted proteome. Out of the 480 proteins predicted to be encoded by *M. kamchatka* 463 had their closest homologs in *M. sibericum*, with 92% identical residues in average. After clustering the paralogs, these proteins corresponded to 434 distinct genes clusters delineating a first estimate of the mollivirus core gene set. Four hundred and eleven of these clusters contained a single copy (singletons) gene for each strain. Remarkably, 290 of the 480 (60.4 %) *M. kamchatka*-encoded proteins did not exhibit a detectable homolog among cellular organisms or previously sequenced viruses (excluding *M. sibericum*). Those will be referred to as “ORFans”. Among the 190 proteins exhibiting significant (E<10^−5^) matches in addition to their *M. sibericum* counterparts, 78 (16% of the total gene content) were most similar to Pandoravirus predicted proteins, 18 (3.7%) to proteins of other virus families, 51 (10.6%) to *A. castellanii* proteins, 24 (5%) to proteins of other eukaryotes, 17 (3.5%) to bacterial proteins and 2 (0.4%) to proteins of *Archaea* (Fig. 3). The interpretation of these statistics are ambiguous as, on one hand, the large proportion of “ORFans” (>60%) is characteristic of what is usually found for the prototypes of novel giant virus families (7). On the other hand, the closest viral homologs are not scattered in diverse previously defined virus families, but mostly belongs to the Pandoraviridae (78/96=81%) (Fig. 3). The two molliviruses thus constitute a new group of viruses with their own specificity but with a phylogenetic affinity with the pandoraviruses, as previously noticed (7). The proportion of *M. kamchatka* proteins with best matching counterparts in Acanthamoeba confirms the high gene exchange propensity with the host, already noticed for *M. sibericum* (6).

**Fig 3.**
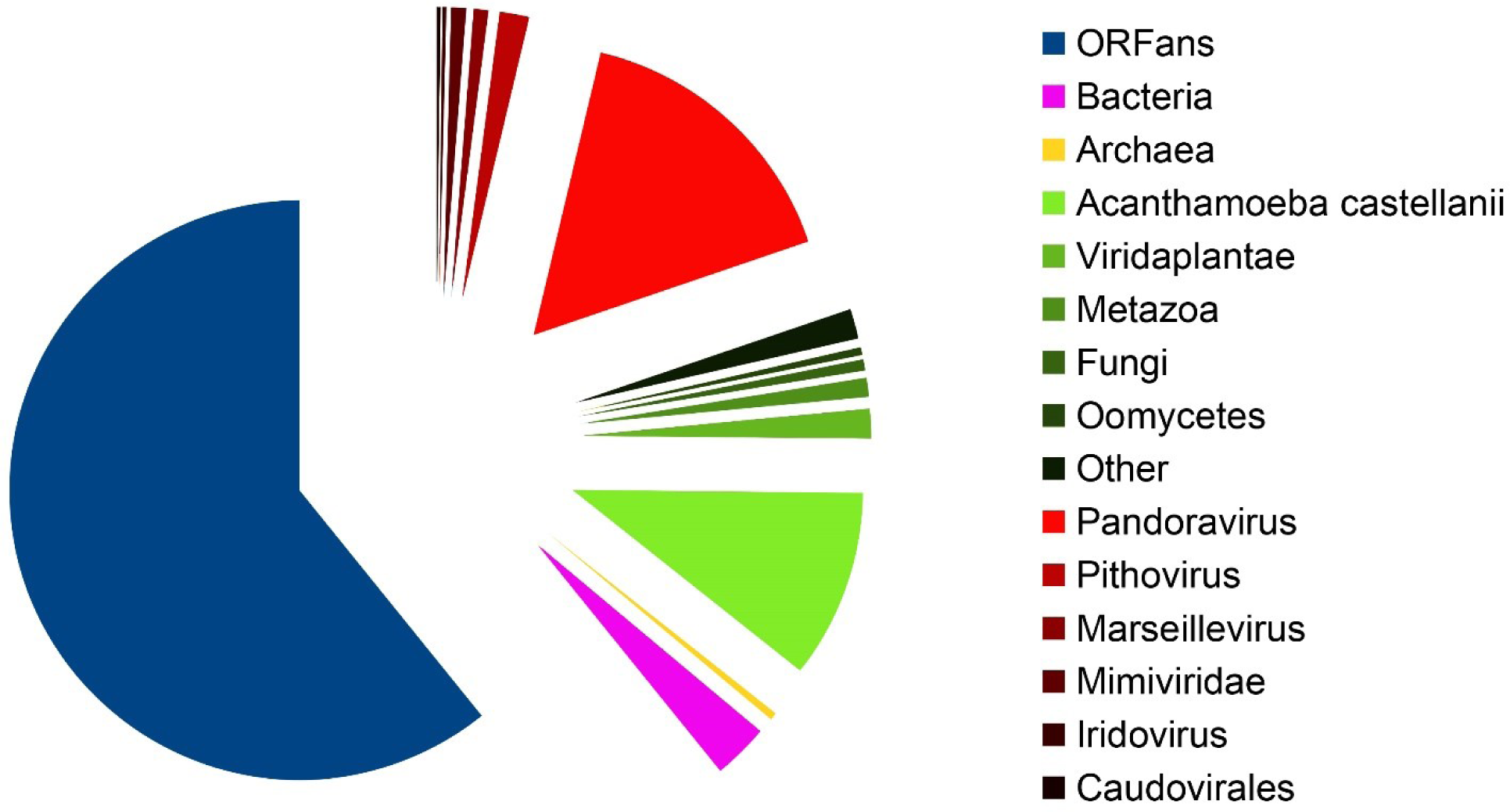
Distribution of the best-matching NR homologs of *M. kamchatka* predicted proteins. Best-matching homologous proteins were identified using BLASTP (E value <10^−5^) against the non-redundant (NR) database (15) (after excluding *M. sibericum*). Green shades are used for eukaryotes, red shades for viruses.

### Recent evolutionary events since the *M. sibericum*/*M. kamchatka* divergence

We investigated the evolutionary events specific of each of the molliviruses by focusing on proteins lacking reciprocal best matches between the two strains. We found 63 such cases of which 10 corresponded to unilateral strain-specific duplications of genes, and 53 were unique to a given strain. These unique genes (Table 1 & 2) result from gains or losses in either of mollivirus strains (20 in *M. Kamchatka*, 33 in *M. sibericum*). The likely origins of these strain-specific genes (horizontal acquisition, *de novo* creation (13, 14), or differential loss) are listed in Table 1 and Table 2.

**Table 1.** Status of the protein-coding genes unique to *M. kamchatka*.

| ORF ID | Predicted function | Putative evolutionary scenario |
| --- | --- | --- |
| mk_25 | None (ORFan) | *de novo* creation or loss in *M. sibericum* |
| mk_92 | DNA methyltransferase | loss in *M. sibericum* (present in some pandoraviruses) |
| mk_93 | Ring domain | loss in *M. sibericum* (present in some pandoraviruses) |
| mk_104 | Ring domain | loss in *M. sibericum* (present in some pandoraviruses) |
| mk_127 | None (ORFan) | *de novo* creation or loss in *M. sibericum* |
| mk_159 | None | loss in *M. sibericum* (present in some pandoraviruses) |
| mk_165 | None | HGT from Pandoravirus |
| mk_166 | Peptidase | loss in *M. sibericum* (present in some pandoraviruses) |
| mk_172 | None (ORFan) | *de novo* creation or loss in *M. sibericum* |
| mk_182 | None (ORFan) | *de novo* creation or loss in *M. sibericum* |
| mk_231 | None (ORFan) | *de novo* creation or loss in *M. sibericum* |
| mk_313 | None (ORFan) | *de novo* creation or loss in *M. sibericum* |
| mk_369 | None (ORFan) | *de novo* creation or loss in *M. sibericum* |
| mk_415 | None (ORFan) | *de novo* creation or loss in *M. sibericum* |
| mk_441 | None (ORFan) | *de novo* creation or loss in *M. sibericum* |
| mk_466 | None (ORFan) | *de novo* creation or loss in *M. sibericum* |
| mk_467 | None | loss in *M. sibericum* (HGT to Acanthamoeba) |
| mk_469 | B1-1 like | loss in *M. sibericum* (present in Acanthamoeba) |
| mk_476 | None (ORFan) | *de novo* creation or loss in *M. sibericum* |
| mk_478 | None (ORFan) | *de novo* creation or loss in *M. sibericum* |

**Table 2.** Status of the protein-coding genes unique to *M. sibericum*.

| ORF ID | Predicted function | Putative evolutionary scenario |
| --- | --- | --- |
| ms_1 | None (ORFan) | *De novo* creation or loss in *M. kamchatka* |
| ms_3 | None (ORFan) | *De novo* creation or loss in *M. kamchatka* |
| ms_5 | None (ORFan) | *De novo* creation or loss in *M. kamchatka* |
| ms_7 | None (ORFan) | *De novo* creation or loss in *M. kamchatka* |
| ms_8 | None (ORFan) | *De novo* creation or loss in *M. kamchatka* |
| ms_13 | None (ORFan) | *De novo* creation or loss in *M. kamchatka* |
| ms_14 | None | HGT to *Acanthamoeba* |
| ms_38 | None (ORFan) | *De novo* creation or loss in *M. kamchatka* |
| ms_42 | None (ORFan) | *De novo* creation or loss in *M. kamchatka* |
| ms_53 | None (ORFan) | *De novo* creation or loss in *M. kamchatka* |
| ms_64 | None | loss in *M. kamchatka* (present in *Noumeavirus*) |
| ms_109 | None (ORFan) | *De novo* creation or loss in *M. kamchatka* |
| ms_120 | None (ORFan) | *De novo* creation or loss in *M. kamchatka* |
| ms_136 | None (ORFan) | *De novo* creation or loss in *M. kamchatka* |
| ms_138 | None (ORFan) | *De novo* creation or loss in *M. kamchatka* |
| ms_139 | None (ORFan) | *De novo* creation or loss in *M. kamchatka* |
| ms_144 | None (ORFan) | *De novo* creation or loss in *M. kamchatka* |
| ms_157 | None (ORFan) | *De novo* creation or loss in *M. kamchatka* |
| ms_159 | None (ORFan) | *De novo* creation or loss in *M. kamchatka* |
| ms_166 | None (ORFan) | *De novo* creation or loss in *M. kamchatka* |
| ms_172 | None (ORFan) | *De novo* creation or loss in *M. kamchatka* |
| ms_190 | None | loss in *M. kamchatka* (present in some pandoraviruses) |
| ms_193 | None (ORFan) | *De novo* creation or loss in *M. kamchatka* |
| ms_246 | None (ORFan) | *De novo* creation or loss in *M. kamchatka* |
| ms_258 | None (ORFan) | *De novo* creation or loss in *M. kamchatka* |
| ms_311 | Zinc-finger domain | loss in *M. kamchatka* (present in *Gossypium hirsutum*) |
| ms_312 | Zinc-finger domain | loss in *M. kamchatka* (present in *Cavenderia fasciculata*) |
| ms_313 | None | loss in *M. kamchatka* (present in some pandoraviruses) |
| ms_464 | None (ORFan) | *De novo* creation or loss in *M. kamchatka* |
| ms_465 | None (ORFan) | *De novo* creation or loss in *M. kamchatka* |
| ms_479 | DNA methyltransferase | loss in *M. kamchatka* (present in some pandoraviruses) |
| ms_494 | None (ORFan) | *De novo* creation or loss in *M. kamchatka* |
| ms_495 | None (ORFan) | *De novo* creation or loss in *M. kamchatka* |

Six *M. kamchatka* proteins, absent from *M. sibericum*, have homologs in pandoraviruses suggesting common gene ancestors (and loss in *M. sibericum*) or horizontal acquisitions. According to its embedded position within the pandoravirus phylogenetic tree, only one anonymous protein (mk_165) could be interpreted as a probable horizontal transfer from pandoraviruses (Fig. 4A). Another candidate, mk_92, shares 75% of identical residues with a pandoravirus DNA methyltransferase (pqer_cds_559). However the very long branch associated to the P. dulcis homolog (eventually due to a non-orthologous replacement) raises some doubt as for the origin of the *M. kamchatka* gene (Fig. 4B).

**Fig 4.**
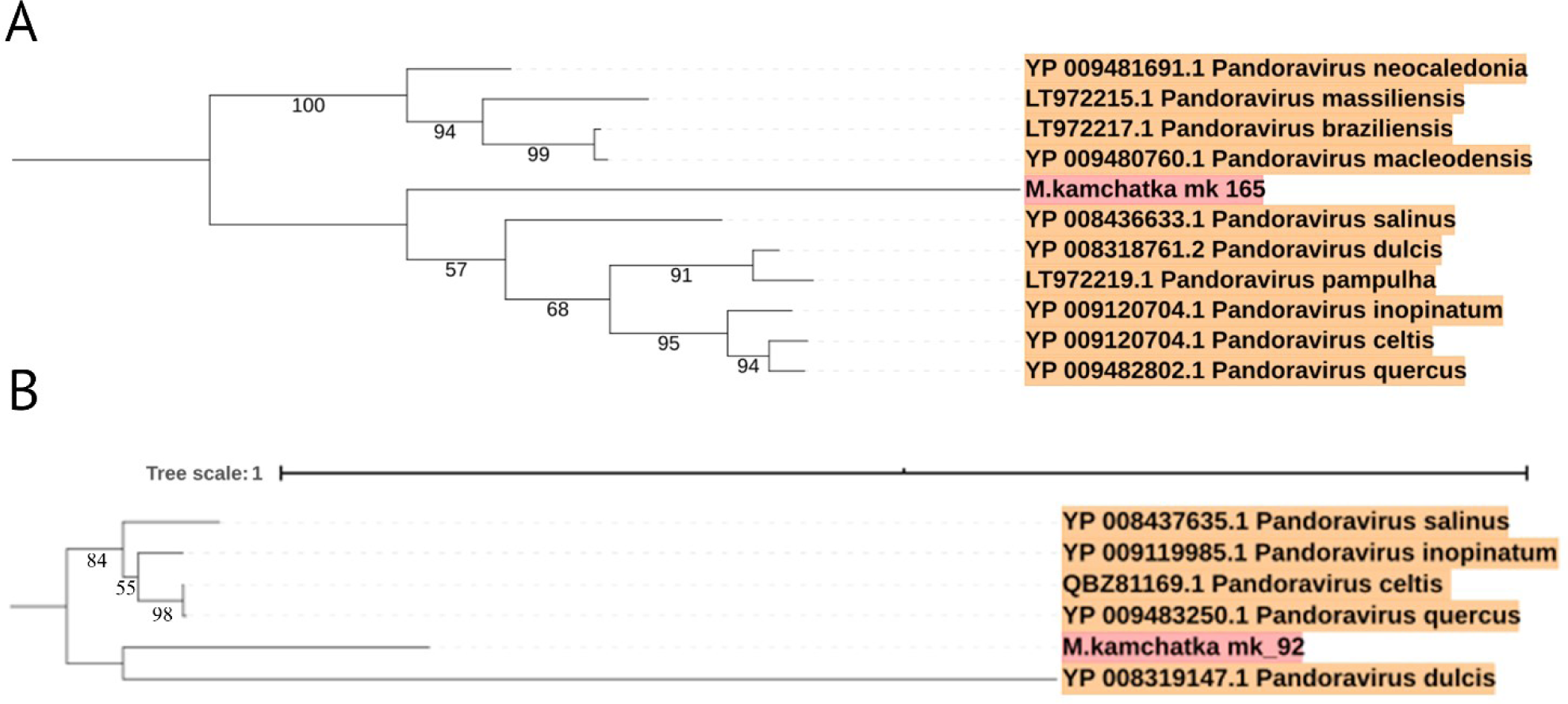
Eventual gene transfers from a pandoravirus to *M. kamchatka*. Both phylogenetic trees were computed from the global alignments of orthologous protein sequences using MAFFT (29). IQtree (36) was used to determine the optimal substitution model (options: « -m TEST » and « –bb 1000 »). (A) Protein mk_165 (no predicted function). (B) Predicted methyltransferase mk_92. Both *M. kamchatka* protein sequences appear embedded within the pandoravirus trees. In (A), the long branch leading to the *M. kamchatka* homolog suggests its accelerated divergence after an ancient acquisition from a pandoravirus. In (B), the long branch leading to the *P. dulcis* homolog might alternatively be interpreted as a non-orthologous replacement of the ancestral pandoravirus version of the gene.

Two *M. kamchatka*-specific proteins, encoded by adjacent genes (mk_466, mk_467), have homologs in Acanthamoeba, suggesting potential host-virus exchanges. Phylogenetic reconstruction did not suggest a direction for the transfer of mk_466. However, since the unique homolog of mk_467 is found in Acanthamoeba (and not in other eukaryotes), the corresponding gene probably originated from a close relative of *M. kamchatka* and was recently transferred to its host.

Three proteins unique to *M. sibericum* have homologs in pandoraviruses (Table 2), suggesting common gene ancestors (and loss in *M. kamchatka*) or horizontal acquisitions. One protein (ms_14) has a unique homolog in Acanthamoeba, suggesting a virus to host exchange. The homolog of ms_312 in *Cavenderia fasciculata* (16) might be the testimony of past interactions between molliviruses and deeply rooted ancestor of the Amoebozoa clade.

The above analyses of the genes unique to each mollivirus indicate that if horizontal transfer may contribute to their presence, it is not the predominant mechanisms for their acquisition. We then further investigated the 12 genes unique to *M. kamchatka* and 26 genes unique to *M. sibericum* (i.e. “strain ORFans” w/o homolog in the databases) by computing three independent sequence properties (Fig. 5): the codon adaptation usage index (CAI), the G+C composition, and the ORF length.

**Fig 5.**
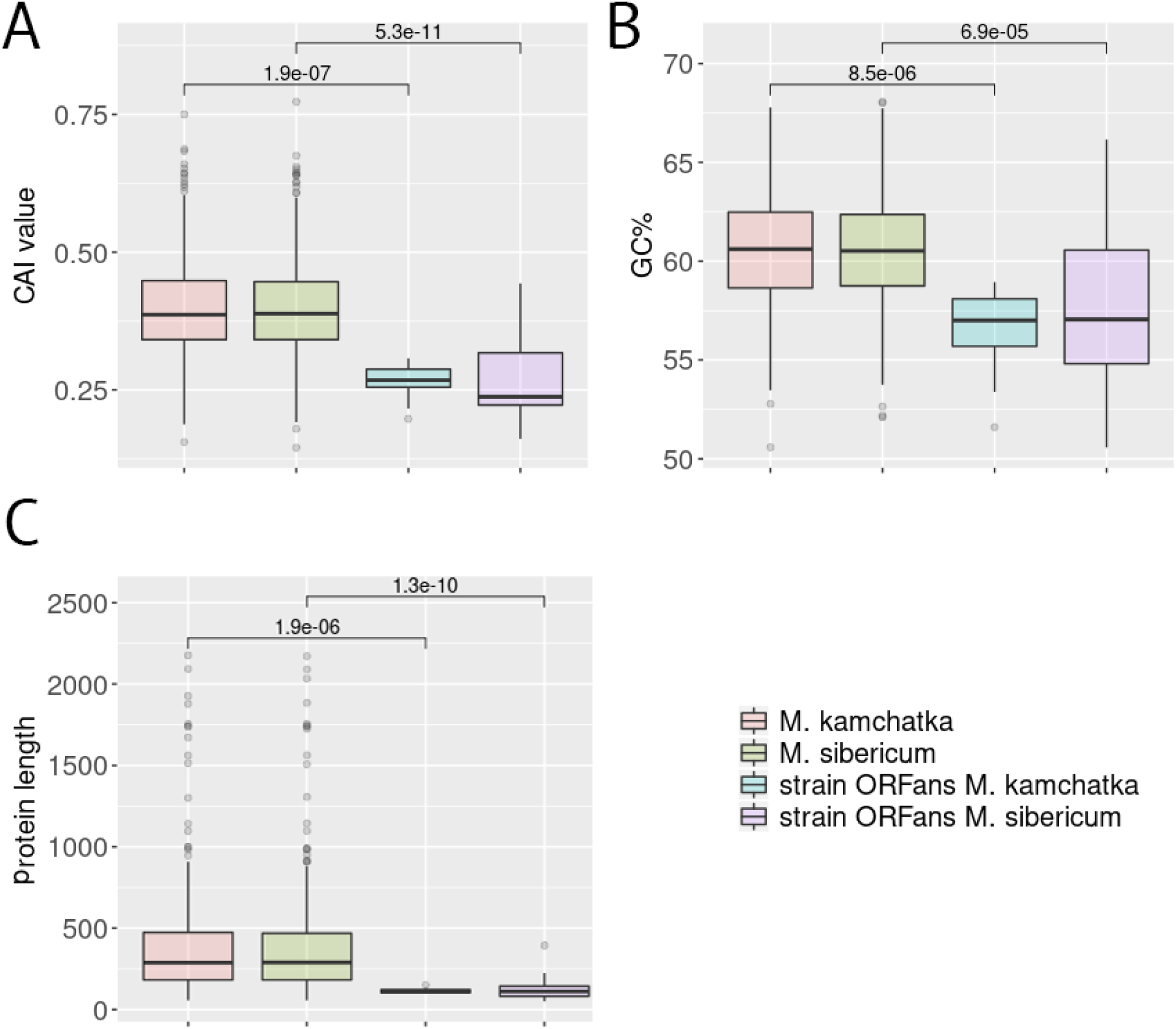
Genomic features of strain-specific ORFans. (A) Codon adaption index (CAI). (B) G+C content. (C) Protein length. Box plots show the median, the 25th and 75th percentiles. P-value are calculated using the Wilcoxon test.

With an average CAI value of 0.26, the strain-specific ORFans appear significantly different from the rest of the mollivirus genes (mean = 0.40, Wilcoxon test p< 2 10^−7^). These genes also exhibit a significantly lower G+C content (56% for *M. kamchatka* and 57% for *M. sibericum*) than the rest of the genes (60.5% for both viruses), also closer to the value computed for intergenic regions (54% in average for both viruses). Moreover, the strain-specific ORFans are smaller in average compared to the rest of the genes (115bp/378bp for *M. kamchatka* and 122bp/369bp for *M. sibericum*). Altogether, those results suggest that *de novo* gene creation might occur in the intergenic regions of molliviruses as already postulated for pandoraviruses (13, 14).

### New predicted protein functions in *M. kamchatka*

Sixty four of the *M. kamchatka* predicted proteins exhibit sequence motifs associated to known functions. Fifty nine of them are orthologous to previously annotated genes in *M. sibericum* (6). This common subset confirms the limited complement of DNA processing and repair enzymes found in molliviruses: mainly a DNA polymerase: mk_287, a primase: mk_236, and 3 helicases: mk_291, mk_293, mk_351. *M. kamchatka* confirms the absence of key deoxynucleotide synthesis pathways (such as thymidylate synthase, thymidine kinase and thymidylate kinase), and of a ribonucleoside-diphosphate reductase (present in pandoraviruses). The five *M. kamchatka* specific proteins (Table 1) associated to functional motifs or domain signature correspond to:

- two proteins (mk_93 and mk_104) containing a type of zinc finger (Ring domain) mediating protein interactions,
- one protein (mk_469) with similarity to the (BI)-1 like family of small transmembrane proteins,
- one predicted LexA-related signal peptidase (mk_166),
- one DNA methyltransferase (mk_92).

### Evaluation of the selection pressure exerted on mollivirus genes

The availability of two distinct strains of mollivirus allows the first estimation of the selection pressure exerted on their shared genes during their evolution. This was done by computing the ratio ω=dN/dS of the rate of non-synonymous mutations (dN) over the rate of synonymous mutations (dS) for pairs of orthologous genes. ω values much lesser than one are associated to genes the mutation of which have the strongest negative impact on the virus fitness. The high sequence similarity of proteins shared by *M. sibericum* and *M. kamchatka* allowed the generation of flawless pairwise alignments and the computation of highly reliable ω values for most (*i.e.* 397/411) of their orthologous singletons.

Fourteen singleton pairs were not taken into account in the selection pressure analysis because of their either identical or quasi-identical sequences (11 of them), or unreliable pairwise alignments (3 of them). For the 397 gene pairs retained in the analysis, the mean ω value was 0.24 ± 0.14 (Fig. 6). This result corresponds to a strong negative selection pressure indicating that most of the encoded proteins greatly contribute to the molliviruses’ fitness. Together with the high level of pairwise similarity (92%) of their proteins, this also indicates that *M. kamchatka* evolved very little during the last 30,000 years and that the *M. sibericum* genome was not prominently damaged during its cryostasis in permafrost.

**Fig 6.**
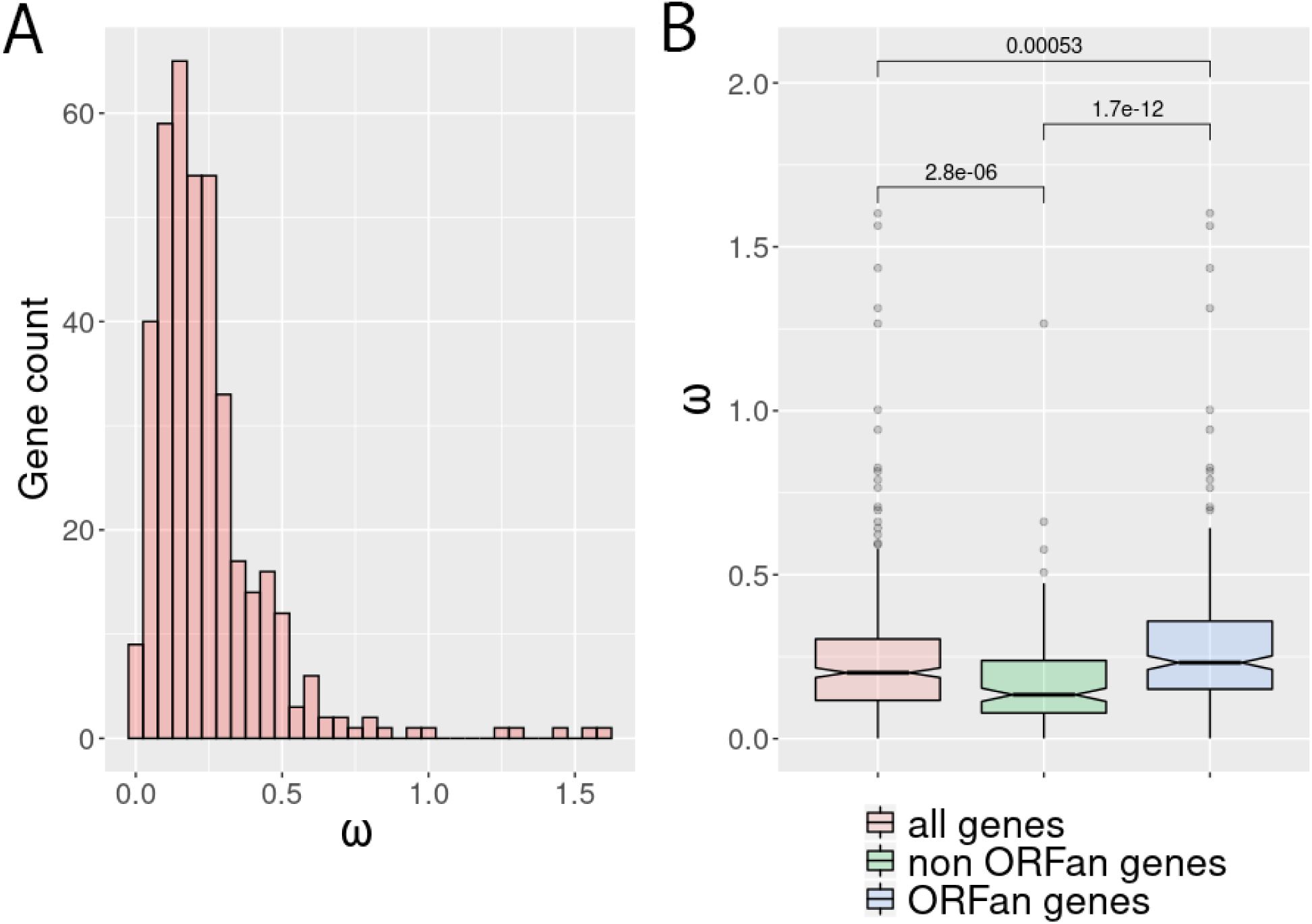
Selection pressure among different classes of genes. Values of ω (i.e. dN/dS) were computed from the alignments of homologous coding regions in *M. kamchatka* and *M. sibericum*. (A) Distribution of calculated ω values (n=397). (B) Box plots of the ω ratio among ORFan genes (n=243) and non ORFan genes (n=154). Box plots show the median, the 25th, and 75th percentiles. All p-value are calculated using the Wilcoxon test.

The analysis restricted to the 244 pairs of ORFan-coding genes resulted in a very similar ω value of 0.29± 0.15 (Fig. 6). This indicates that although homologs of these proteins are only found in molliviruses, they have the same impact on the virus fitness than more ubiquitous proteins. This confirms that they do encode actual proteins, albeit with unknown functions. In contrast, four orthologous pairs (ms_160/mk_141; ms_280/mk_262; ms_171/mk_151; ms_430/mk_411; ms_60/mk_48) exhibit ω value larger than one. Those ORFans either are under positive selection for maintaining their functions, or newly created gene products undergoing refinement or pseudogenization.

We further examined the selection pressure of proteins-coding genes with homologs in pandoraviruses. We used their 10 sequenced genomes to generate the corresponding gene clusters (Fig. 7). The 90 clusters shared by both virus groups included 64 singletons (single copy gene present in all viruses), among which 55 were suitable for dN/dS computations. The mean ω value (0.17 ± 0.1) was very low, indicating that these genes, forming a “super core” gene set common to the molliviruses and pandoraviruses, are under an even stronger negative selection pressure than those constituting the provisional (most likely overestimated) mollivirus core gene set.

**Fig 7.**
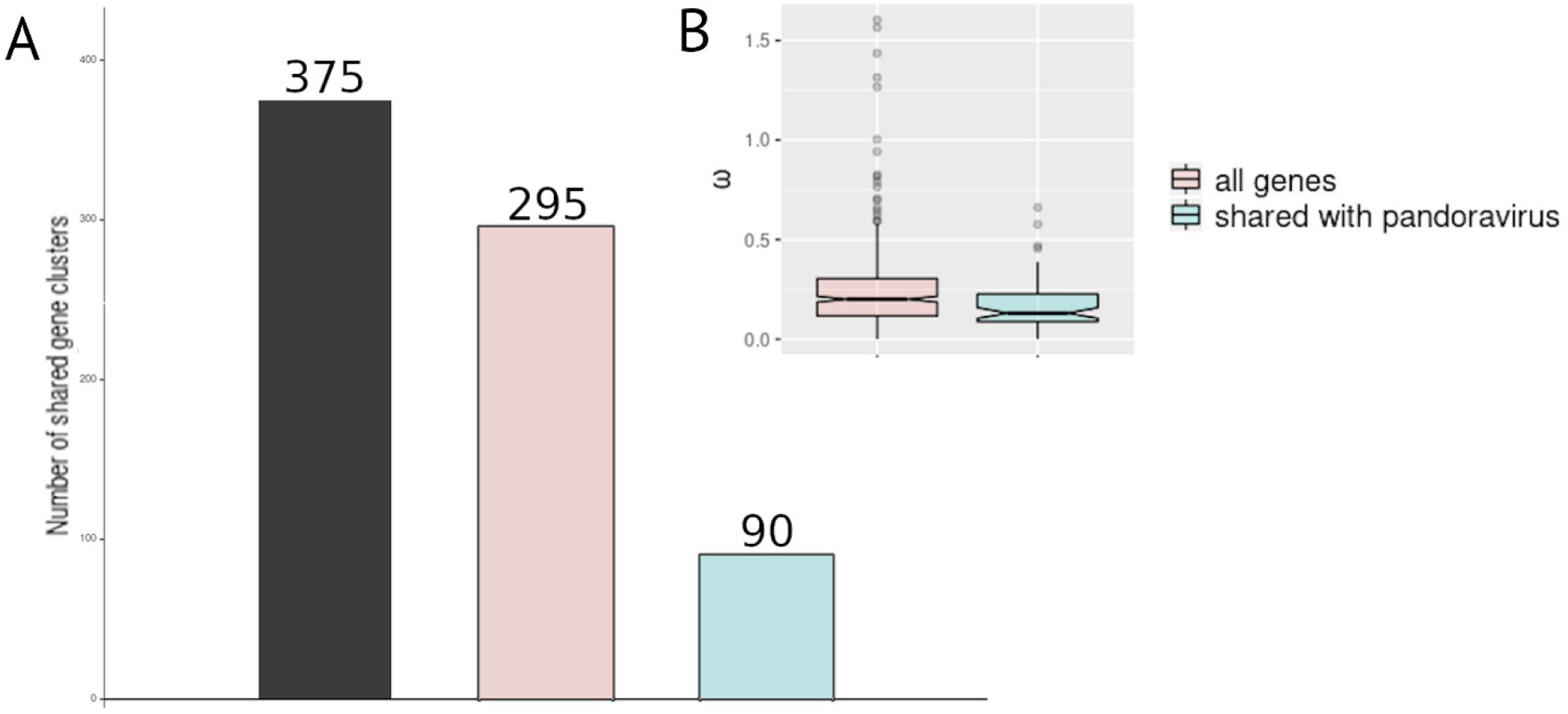
Comparison of the mollivirus and pandoravirus core gene contents. (A) The distribution of the protein clusters shared by all pandoraviruses (black), the two Molliviruses (pink), and by both virus groups (super core genes) (blue). (B) Box plot of ω values calculated from the alignment of molliviruses core genes (pink), and super core genes (blue). Box plots show the median, the 25th, and 75th percentiles.

### Genomic inhomogeneity

The original genome analysis of Lausannevirus (a member of the Marseilleviridae family) (17) revealed an unexpected non-uniform distribution of genes according to their annotation. “Hypothetical” genes (i.e. mostly ORFans) were segregated from “annotated” (i.e. mostly non-ORFans) in two different halves of the genome. In a more recent work, we noticed a similar bias in the distribution of Pandoravirus core genes (13). The availability of a second mollivirus isolate gave us the opportunity to investigate this puzzling feature for yet another group of Acantamoeba-infecting virus. In Fig. 8, we plotted the distribution of three types of genes: 1-those with homologs in *A. castellanii* (n= 55 for *M. sibericum* and n= 51 for *M. kamchatka*), 2-those belonging to the super core set shared by both molliviruses and pandoraviruses (n=64), 3-those unique to either mollivirus strains (n=26 for *M. sibericum*, n=12 for *M. kamchatka*). These plots reveal a strong bias in the distribution of the super core vs. ORFan genes (Fig. 8). The first half of the *M. sibericum* genome exhibits 90% of its ORFans while the second half contains most of the members of the super core gene set. In contrast, genes eventually exchanged with the host display a more uniform distribution. The lack of an apparent segregation in the distribution of ORFans in the *M. kamchatka* genome might be due to their underprediction as no transcriptome information is available for this strain. Fig. 9 shows that there is also a strong bias in the distribution of single-copy genes *vs.* those with paralogs in either *M. sibericum* and/or *M. kamchatka*. Altogether, these analyses suggest that the two genome halves follow different evolutionary scenario, the first half concentrating the genomic plasticity (*de novo* gene creation, gene duplication), the other half concentrating the most conserved, eventually essential, gene content.

**Fig 8.**
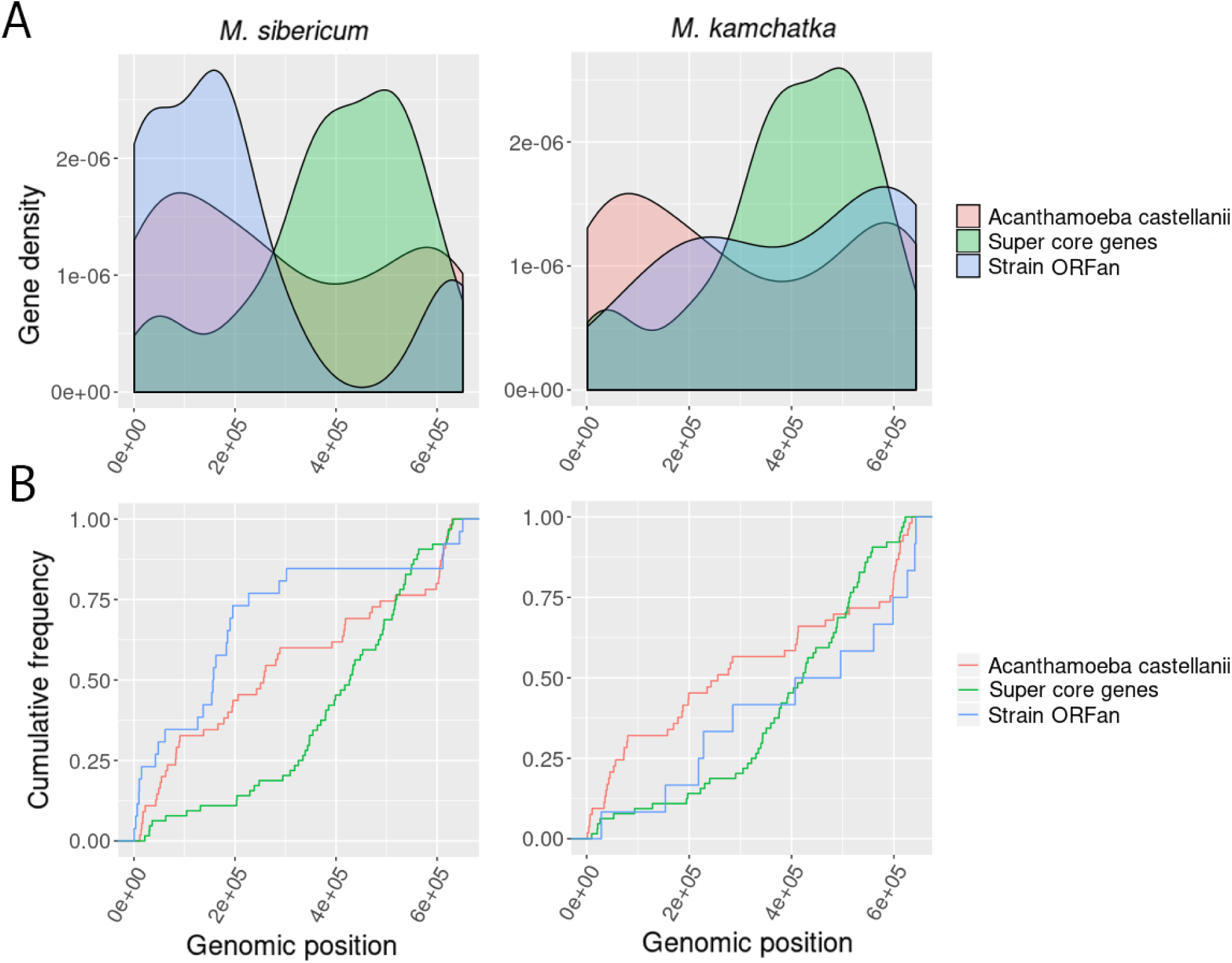
Distribution of different classes of genes along mollivirus genomes. (A) Variation of the gene density as computed by the ggpplot2 “geom_density” function (37). Genes with best-matching homologs in *A. castellanii* (in the NR database excluding mollivirus) are uniformly distributed (in pink) in contrast to super core genes (in green) and strain-specific ORFans (in blue). (B) Cumulative distribution of the above classes of genes using the same color code.

**Fig 9.**
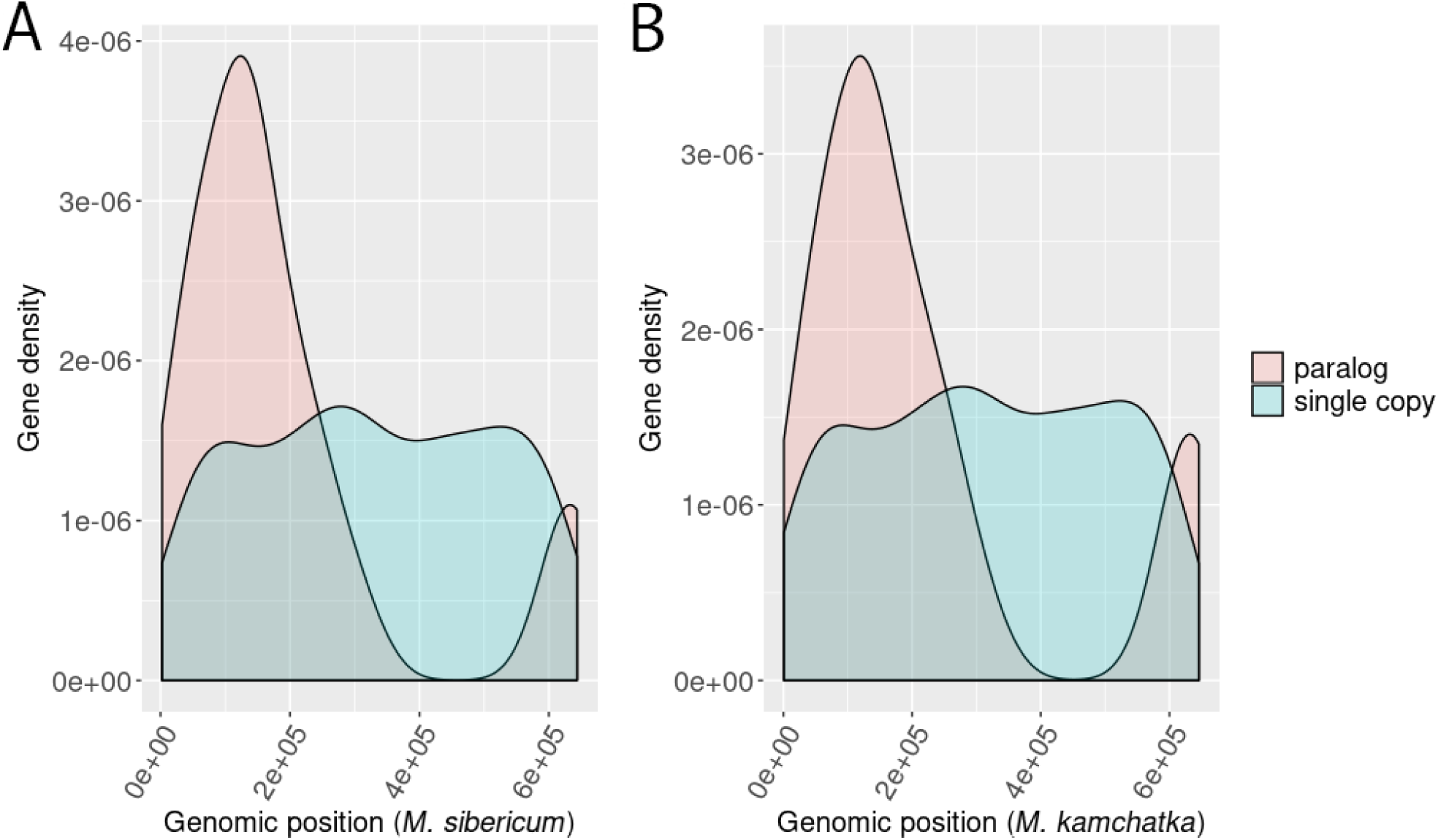
Distribution of single-copy *vs.* multiple copy genes along mollivirus genomes. Single copy genes (in blue) in both strains are uniformly distributed in contrast to genes with paralogs (pink) in at least one strain that cluster in the left half of the genomes. (A) *M. sibericum* (n=48). (B) *M. kamchatka* (n= 46).

## Discussion

Following the discovery of their first representatives, each families of giant (e.g. Mimiviridae, Pithoviridae, Pandoraviridae) and large (e.g. Marseilleviridae) viruses infecting acanthamoeba have expanded steadily, suggesting they were relatively abundant and present in a large variety of environments. One noticeable exception has been the molliviruses, the prototype of which remained unique after its isolation from 30,000-year old permafrost. The absence of *M. sibericum* relatives from the large number of samples processed by others and us since 2014, raised the possibility that they might have gone extinct, or might be restricted to the Siberian arctic. Our isolation of a second representative of the proposed Molliviridae family, *M. kamtchatka*, at a location more than 1,500 km from the first isolate and enjoying a milder climate, is now refuting these hypotheses. Yet, the planet-wide ubiquity of these viruses remains to be established, in contrast to other acanthamoeba-infecting giant viruses (7). Even when present, mollivirus-like viruses appear to be in very low abundance, as judged from the very small fraction of metagenomics reads they represent in total sample DNA for *M. kamchatka* (about 0.02 part per million) as well as for *M. sibericum* (about one part per million)(6). Another possibility would be that the actual environmental host is not an acanthamoeba, the model host used in our laboratory. However, evidences of specific gene exchanges with acanthamoeba (including a highly conserved homolog major capsid protein) (6, 18, 19) make this explanation unlikely. We conclude that members of the proposed Molliviridae family are simply less abundant than other acanthamoeba-infecting viruses, a conclusion further supported by the paucity of Mollivirus-related sequences in the publicly available metagenomics data (data not shown).

As always the case, the characterization of a second representative of a new virus representative opened new opportunities of analysis. Unfortunately, the closeness of *M. kamchatka* with *M. sibericum* limited the amount of information that could be drawn from their comparison. For instance, the number of genes shared by the two isolates is probably a large overestimate of the “core” gene set characterizing the whole family. On the other hand, the closeness of the two isolates allowed an accurate determination of the selection pressure (ω=dN/dS) exerted on many genes, showing that most of them, including mollivirus ORFans, encode actual proteins the sequence of which are under strong negative selection and thus contribute to the virus fitness. Given the partial phylogenetic affinity (i.e. 90 shared gene clusters) of the mollivirus with the pandoraviruses, we also assessed the selection pressure exerted on 55 of these “super core” genes, and found them under even stronger negative selection (Fig. 7). This suggests that this super core gene set might have been present in a common ancestor to both proposed families.

If we postulate that *M. sibericum* underwent into a complete stasis when it became frozen in permafrost while *M. kamchatka* remained in contact with living acanthamoeba, we could consider the two viral genomes to be separated by at least 30,000 years of evolution (eventually more if they are not in a direct ancestry relationship)(20). The high percentage of identical residues (92%) in their proteins corresponds to a low substitution rate of 1.7 10^−6^ amino acid change/position/year. This is an overestimate since the two viruses probably started to diverge from each other longer than 30,000-year ago. This value is nevertheless comparable with estimates computed for poxviruses (21) given the uncertainty on the number of replicative cycles occurring per year. The high level of sequence similarity of *M. kamchatka* with *M. sibericum* also indicates that the later did not suffer much DNA damage during its frozen stasis, even in absence of detectable virus-encoded DNA repair functions.

Horizontal gene transfers with the host were suggested by the fact that 51 proteins shared by the two mollivirus strains exhibited a second best match in acanthamoeba. Because no homolog is detected in other eukaryotes for most of them, these transfers probably occurred in the mollivirus-to-host direction. The clearest case is that of a major capsid protein homolog (mk_314, ml_347) sharing 64% identical residues with a predicted acanthamoeba protein (locus: XP_004333827). Two other genes encoding proteins that have also homologs in molliviruses flank the corresponding host gene. However, the corresponding viral genes are not collinear in *M. sibericum* or *M. kamchatka* and were probably transferred from a different, yet unknown mollivirus strain. The presence of a 100% conserved major capsid protein homolog in the genome of *M. kamchatka* and *M. sibericum* is itself puzzling. Such protein (with a double-jelly roll fold) is central to the structure of icosahedral particles (22). Consistent with its detection in *M. sibericum* virions (6), its conservation in *M. kamchatka*, suggests that it still plays a role in the formation of the spherical mollivirus particles, while it has no homolog in the pandoraviruses. Inspired by previous observations made on the unrelated Lausannevirus genome (17), we unveiled a marked asymmetry in the distribution of different types of protein-coding genes in the Mollivirus genomes. As shown in Fig. 8 the left half of the genome concentrates most of the genes coding for strain-specific ORFans while the right half concentrates most of super core genes shared with pandoraviruses. This asymmetry is even stronger for the multiple copy genes while single-copy genes are uniformly distributed along the genome (Fig. 9). The molliviruses thus appear to confine their genomic “creativity” (*de novo* creation and gene duplication) in one-half of their genome, leaving the other half more stable. An asymmetry in the distribution of the core genes was previously noticed in the pandoravirus genomes (13). Such features might be linked to the mechanism of replication that is probably similar for the two virus families. Further studies are needed to investigate this process. The asymmetrical genomic distribution of pandoravirus core genes and mollivirus super core genes might be a testimony of their past common ancestry.

Despite their differences in morphology, as well as in virion and genome sizes, the comparative analysis of the prototype *M. sibericum* and of the new isolate *M. kamchatka* confirms their phylogenetic affinity with the Pandoraviruses (Fig. 3, Fig. 10). However, it remains unclear whether this is due to a truly ancestral relationship between them, or if it is only the consequence of numerous past gene exchanges favored by the use of the same cellular host. From the perspective of the sole DNA polymerase sequence, the two known molliviruses do cluster with the pandoraviruses, albeit at a larger evolutionary distance than usually observed between members of the same virus family (Fig. 10). In absence of an objective threshold, and pending the characterization of eventual “missing links”, we thus propose to classify *M. sibericum* and *M. kamchatka* as members of the proposed Molliviridae family, distinct from the Pandoraviridae.

**Fig 10.**
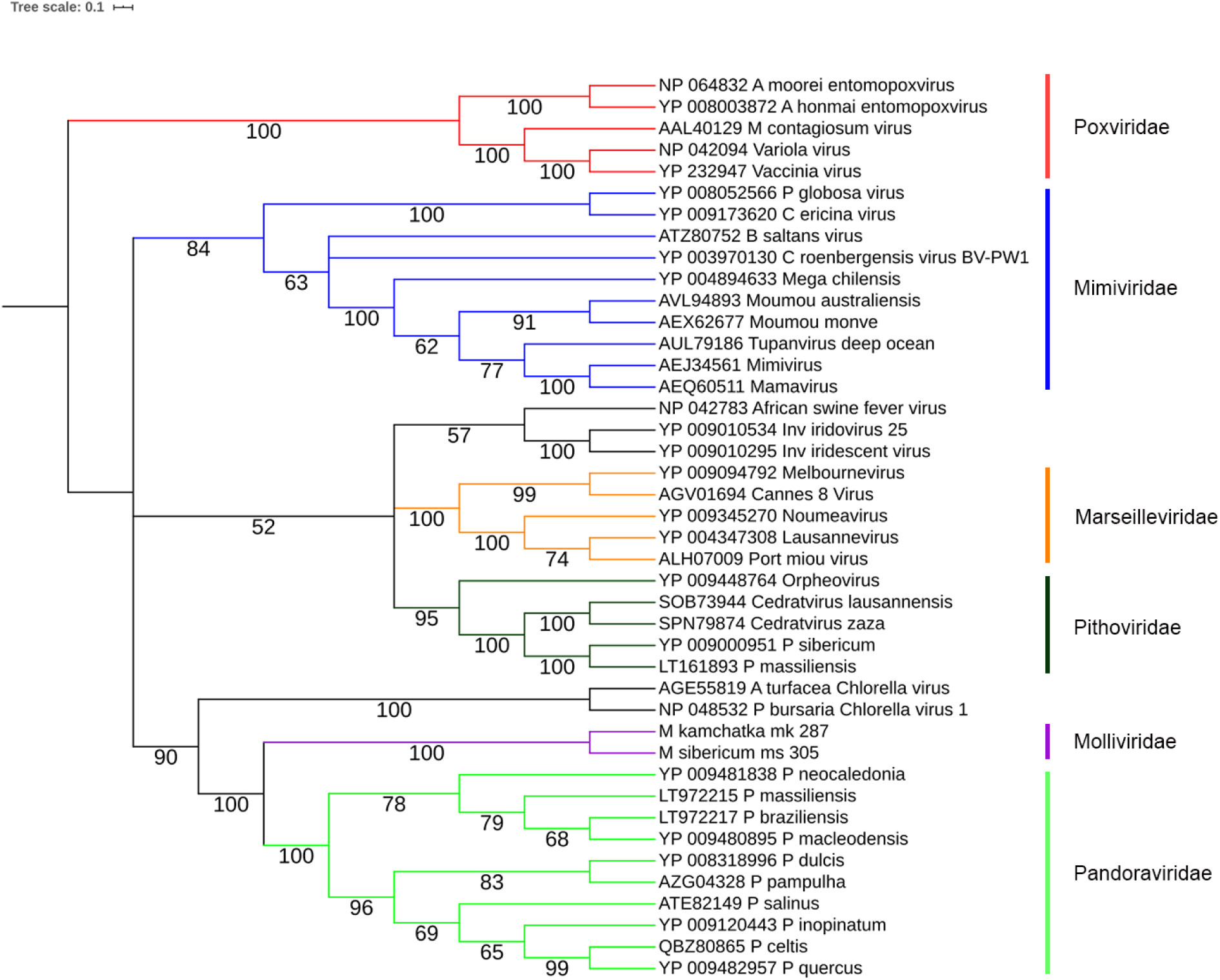
Phylogeny of DNA polymerase B of large and giant dsDNA viruses. This neighbor-joining tree was computed (JTT substitution model, 100 resampling) on 397 amino acid positions from an alignment of 42 sequences computed by MAFFT (29). Branches with boostrap values <60% were collapsed.

## Materials and Methods

### Virus isolation

We isolated *M. kamchatka* from muddy grit collected near Kronotski Lake, Kamchatka (Russian Federation N :54 32 59, E :160 34 55). The sample was stored for twenty days in pure rice medium (23) at room temperature. An aliquot of the pelleted sample triggered an infected phenotype on a culture of *Acanthamoeba castellanii Neff* (ATCC30010TM) cells adapted to 2,5µg/mL of Amphotericin B (Fungizone), Ampicillin (100µg/ml), Chloramphrenicol (30µg/ml) and Kanamycin (25µg/ml) in protease-peptone–yeast-extract–glucose (PPYG) medium after two days of incubation at 32°C. A final volume of 6 mL of supernatant from two T25 flasks exhibiting infectious phenotypes was centrifuged for 1 hour at 16,000xg at room temperature. Two T75 flasks were seeded with 60,000 cells/cm^2^ and infected with the resuspended viral pellet. Infected cells were cultured in the same conditions as described below. We confirmed the presence of viral particles by light microscopy.

### Validation of the presence of *M. kamchatka* in the original sample

To confirm the origin of the *M. kamchatka* isolate from the soil of the Kronotsky river bank, DNA was extracted from the sample and sequenced on an Illumina platform, leading to 340,320,265 pair-ended reads (mean length 150bp). These metagenomics reads were then mapped onto the genome sequence of *M. kamchatka*. Seven matching (100 % Identity) pair-ended reads (hence 14 distinct reads) were detected, indicating the presence of virus particles in the original sample, although at very low concentration. However, the very low probability of such matches by chance (p<10^−63^) together with the scattered distribution of these matches along the viral genome, further demonstrate the presence of *M. kamchatka* in the original sample.

### Virus Cloning

Fresh *A. castellanii* cells were seeded on a 12-well culture plate at a final concentration of 70,000cells/cm^2^. Cell adherence was controlled under light microscopy after 45 minute and viral particles were added to at a multiplicity of infection (MOI) around 50. After 1 h, the well was washed 15 times with 3 mL of PPYG to remove any viral particle in suspension. Cells were then recovered by gently scrapping the well, and a serial dilution was performed in the next three wells by mixing 200μL of the previous well with 500μL of fresh medium. Drops of 0.5μL of the last dilution were recovered and observed by light microscopy to confirm the presence of a unique *A. castellanii* cell. The 0.5μL droplets were then distributed in each well of three 24-well culture plate. Thousand uninfected *A. castellanii* cells in 500μL of PPYG were added to the wells seeded with a single cell and incubated at 32°C until witnessing the evidence of a viral production from the unique clone. The corresponding viral clones were recovered and amplified prior purification, DNA extraction and cell cycle characterization by electron microscopy.

### Virus mass production and purification

A total number of 40 T75 flasks were seeded with fresh *A. castellanii* cells at a final concentration of 60,000 cells/cm^2^. We controlled cell adherence using light microscopy after 45 minute and flasks were infected with a single clone of *M. kamchatka* at MOI=1. After 48h hours of incubation at 32°C, we recovered cells exhibiting infectious phenotypes by gently scrapping the flasks. We centrifuged for 10 min at 500×g to remove any cellular debris and viruses were pelleted by a 1-hour centrifugation at 6,800xg. The viral pellet was then layered on a discontinuous cesium chloride gradient (1,2g/cm^2^/ 1,3g/cm^2^/ 1,4g/cm^2^/ 1,5g/cm^2^) and centrifuged for 20h at 103,000xg. The viral fraction produced a white disk, which was recovered and washed twice in PBS and stored at 4 °C or −80 °C with 7,5% DMSO.

### Infectious cycle observations using TEM

Twelve T25 flasks were seeded with a final concentration of 80,000 cells/cm^2^ in PPYG medium containing antibiotics. In order to get a synchronous infectious cycle eleven flasks were infected by freshly produced *M. kamchatka* at substantial MOI=40. The *A. castellanii* infected flasks were fixed by adding an equal volume of PBS buffer with 5 % glutaraldehyde at different time points after the infection : 1h pi, 2h pi, 3h pi, 4h pi, 5h pi, 6h pi, 7h pi, 8h pi, 9h pi, 10h pi and 25h pi. After 45min of fixation at room temperature, cells were scrapped and pelleted for 5min at 500xg. Then cells were resuspended in 1ml of PBS buffer with 2,5 % glutaraldehyde and stored at 4°C. Each sample was coated in 1mm^3^ of 2 % low melting agarose and embedded in Epon-812 resin. Optimized osmium-thiocarbohydrazide-osmium (OTO) protocol was used for staining the samples: 1h fixation in PBS with 2 % osmium tetroxide and 1.5 % potassium ferrocyanide, 20 min in water with 1 % thiocarbohydrazide, 30min in water with 2 % osmium tetroxide, overnight incubation in water with 1 % uranyl acetate and finally 30min in lead aspartate. Dehydration was made using an increasing concentration of ethanol (50 %, 75 %, 85 %, 95 %, 100 %) and cold dry acetone. Samples were progressively impregnated with an increasing mix of acetone and Epon-812 resin mixed with DDSA 0,34v/v and NMA 0,68v/v (33 %, 50 %, 75 % and 100%). Final molding was made using a hard Epon-812 mix with DDSA 0,34v/v NMA 0,68v/v and 0,031v/v of DMP30 accelerator and hardened in the oven at 60°C for 5 days. Ultrathin sections (90nm thick) were observed using a FEI Tecnai G2 operating at 200kV.

### DNA Extraction

*M. kamchatka* genomic DNA was extracted from approximately 5 × 10^9^ purified virus particles using Purelink Genomic extraction mini kit according to the manufacturer’s recommendation. Lysis was performed with in a buffer provided with the kit and extra DTT at a final concentration of 1mM.

### Genome sequencing and assembly

*M. kamchatka* genome was assembled using Spades (24) with a stringent K-mer parameter using both various iteration steps (k = 21,41,61,81,99,127), the “-careful” option to minimize number of mismatches in the final contigs, and the “-nanopore” option to use long reads.

### Annotation of *Mollivirus sibericum* and *Mollivirus kamchatka*

A stringent gene annotation of *M. sibericum* was performed as previously described (13) using RNA-seq transcriptomic data (6). Stranded RNA-seq reads were used to accurately annotate protein-coding genes. Stringent gene annotation of *M. kamchatka* was performed w/o RNA seq data but taking into account protein similarity with *M. sibericum*. Gene predictions were manually curated using the web-based genomic annotation editing platform Web Apollo (25). Functional annotations of protein-coding genes of both genomes were performed using a two-sided approach as already previously described (13). Briefly, protein domains were searched with the CD-search tool (26) and protein sequence searching based on the pairwise alignment of hidden Markov models (HMM) was performed against the Uniclust30 database using HHblits tool (27). Gene clustering was done using Orthofinder’s default parameters (28) adding the “-M msa -oa” option. Strict orthology between pairs of proteins was confirmed using best reciprocal blastp matches.

### Selection Pressure Analysis

Ratios of non-synonymous (dN) over synonymous (dS) mutation rates for pairs of orthologous genes were computed from MAFFT global alignment (29) using the PAML package and codeml with the « model = 2 » (30). A strict filter was applyied to the dN/dS ratio: dN > 0, dS > 0, dS ≤ 2 and dN/dS ≤ 10. The computation of the Codon Adaptation Index (CAI) of both Mollivirus was performed using the cai tool from the Emboss package (31).

### Metagenome sequencing, assembly and annotation

All metagenomic sample were sequenced with DNA-seq paired-end protocol on Illumina HiSeq platform at Genoscope producing 16 datasets of 2×150bp read length. Raw reads quality was evaluated with FASTQC (32). Identified contaminants were removed and remaining reads were trimmed on the right end using 30 as quality threshold with BBTools (33). Assemblies of filtered data sets were performed using MEGAHIT (34) with the following options: “--k-list 33,55,77,99,127 --min-contig-len 1000”. All filtered reads were then mapped to the generated contigs using Bowtie2 (35) with the “--very-sensitive” option.

## Availability of data

The *M. kamtchatka* annotated genome sequence is freely available from the public through the Genbank repository (URL://www.ncbi.nlm.nih.gov/genbank/) under accession number XXXXX.

## Acknowledgements

We are deeply indebted to our volunteer collaborator Alexander Morawitz for collecting the Kamchatka soil sample. We thank N. Brouilly, F. Richard and A. Aouane (imagery platform, Institut de Biologie du Développement de Marseille Luminy) for their expert assistance. E Christo-Foroux is the recipient of a DGA-MRIS scholarship (201760003). This project has received funding from the European Research Council (ERC) under the European Union’s Horizon 2020 research and innovation program (grant agreement No 832601) and from the FRM prize “Lucien Tartois” to C. Abergel. The funding bodies had no role in the design of the study, analysis, and interpretation of data and in writing the manuscript.

